# Effect of Over-expression of GRXs on Thermo and Acetic Acid Stress Tolerance of *Saccharomyces cerevisiae*

**DOI:** 10.1101/2024.06.24.600531

**Authors:** Lin Wang, Li-shuang Zhang, Mei-ling Zhang, Ya-xin He, Yuan Yu, Ke Xu

**Author notes:** Corresponding author e-mail addresses (Ke Xu).

## Abstract

Ethanol production from renewable cellulosic materials is a globally significant research area. However, the high temperatures and acetic acid generated during cellulose pretreatment can inhibit *Saccharomyces cerevisiae* growth, reducing ethanol yields. This study investigates the impact of glutaredoxin family genes (GRXs) over-expression on *S. cerevisiae* cell growth and fermentation performance under thermal and acetic acid stress. Engineered strains overexpressing *GRX1, GRX2*, and *GRX5* demonstrated enhanced growth at 42°C, while those overexpressing *GRX1, GRX2, GRX6*, and *GRX7* showed improved growth at 1 g/L acetic acid. These results suggest that GRX over-expression can remediate *S. cerevisiae*, potentially accelerating advancements in green biomanufacturing.

## 1. Introduction

Green biomanufacturing has emerged as a sustainable and energy-efficient approach integrated with industrial biotechnology, focusing on the clean production of biofuels and biochemicals [1]. By employing chemical engineering and biotechnological methods, we can harness renewable raw materials and biomass to produce valuable products. Transitioning from traditional chemical manufacturing to green biomanufacturing can mitigate issues such as high emissions, energy consumption, and reliance on fossil fuels for compound synthesis [2]. The development of safe, renewable, and sustainable biomanufacturing systems using renewable biomaterials has garnered global interest for the biosynthesis of valuable products [3]

The bioeconomy is currently shifting its focus from “food biomass” to “non-food biomass,” with cellulosic materials being the most abundant and cost-effective renewable resources available [1]. These include a variety of grass family crops, forestry waste, and fiber waste. Utilizing cellulose through microbial fermentation technology to produce fuel ethanol presents a viable method for generating clean energy and reducing environmental pollution [4]. The performance of brewing yeast, a key component of bioethanol fermentation, is crucial for determining the economic and environmental sustainability of the production process. Ingledew et al. [5] identified “warning levels” for factors affecting yeast growth and fermentation in 2004, noting that exceeding these levels can significantly impact yeast performance and result in negative synergistic effects when multiple stress factors are combined.

Toxic byproducts, such as weak acids, aldehydes, and phenols, can be generated during the pretreatment of lignocellulosic biomass, posing a threat to cell growth and metabolism [6]. Acetic acid, a primary toxic byproduct, can acidify the intracellular environment and significantly inhibit yeast cell growth and metabolism [8]. Its toxicity is linked to both anion inhibition effects dependent on pKa and the damage caused by low pH to the cell [9].

In the process of synchronous saccharification and fermentation with cellulose raw materials, a temperature discrepancy exists between the optimal temperatures for cellulase (around 55°C) and yeast growth (around 30°C) [10]. The fermentation tank”s temperature is typically maintained at 32-35°C due to external factors and the heat produced by yeast metabolism, which hinders efficient saccharification and fermentation [3]. Elevated temperatures can damage yeast cell structures and increase intracellular reactive oxygen species (ROS), leading to oxidative toxicity [11]. Enhancing yeast tolerance to high temperatures can reduce the need for cooling water, save on production costs, and improve the efficiency and economic viability of cellulosic ethanol production.

Common strategies to improve S. cerevisiae”s tolerance to environmental stress include evolutionary engineering, genome shuffling, and global transcriptional regulation mechanisms [12]. Advances in functional genomics, transcriptomics, and metabolomics offer new avenues for strain transformation, with synthetic biology providing a framework for designing and constructing artificial biological systems at the molecular level [13, 14]. Our previous study demonstrated that overexpressing GRX5 significantly enhances S. cerevisiae”s thermo tolerance, suggesting a role for glutaredoxins (GRXs) in yeast”s high-temperature tolerance [16].

GRXs, as thiol oxidoreductases, regulate the redox state of protein disulfides or glutathione-protein mixed disulfides using reduced glutathione (GSH). Under normal conditions, GRXs facilitate protein reduction, but under oxidative stress, they can catalyze the glutathionylation of protein sulfhydryl groups, modulating protein activity [17, 18]. In S. cerevisiae, GRXs are categorized into two types: dimercaptoredoxins with roles in cell redox and monothio glutaredoxins involved in iron metabolism [19]. This study involved the expression of all S. cerevisiae GRXs and validated their effects on fermentation performance under high-temperature and acetic acid stress. Engineered strains overexpressing GRX1, GRX2, and GRX5 showed improved growth at 42°C, while those overexpressing GRX1, GRX2, GRX6, and GRX7 demonstrated enhanced growth at 1 g/L acetic acid. These results suggest that GRX overexpression can remediate S. cerevisiae, potentially accelerating the development of green biomanufacturing.

## 2. Material and methods

### 2.1. Strains, vectors and media

*S. cerevisiae* strain BY4741 (MATa ura3Δ0 leu2Δ0 his3Δ1 met15Δ0,SY022) (Invitrogen, Carlsbad, CA) was used in our research. Engineered strains were cultured at 30°C in SD medium lacking uracil with 20 g/L glucose. *E. coli* Top10 (Novagen, USA) competent cells were used for transformation and plasmid DNA extraction, and were cultivated at 37°C in LB medium with 100 mg/L ampicillin or 100 mg/L kanamycin. The gene-accepting vectors HCKan-P, HCKan-O and HCKan-T for golden gate assembly were provided by Prof. Dai, Shenzhen Institutes of Advanced Technology. Plasmids used in this study are listed in (Supplementary Table 1). The genes and primers were synthesized by GENEWIZ (Suzhou, China).

### 2.2. DNA manipulation

For DNA manipulations, TIAN prep Mini Plasmid Kit (TIANGEN,China) was used to isolate plasmids from *E*.*coli*, respectively. Genomic DNA isolation was performed by using the TIAN amp Yeast DNA Kit (TIANGEN). Enzymes used for recombinant DNA cloning and Golden Gate Assembly were purchased from Thermo Scientific (Waltham, MA) and New England Biolabs (Ipswich, MA). All these genes were cloned from S. cerevisiae and followed by DNA amplification using their respective primers by PCR. PCR products were purified by TIAN quick Midi Purification Kit (TIANGEN) and genetic circuits were constructed with standard vector parts by employing the Golden Gate Assembly[4]. The candidate GRXs genes were ligated into the POT vectors along with *TDH3p* promotor upstream and *SLM5t* terminator downstream, followed by transformation into *S. cerevisiae* strain BY4741. Primers used in the DNA assembly are summarized in (Supplementary Table 2)

### 2.3. Real time reverse transcription PCR

The total RNA was extracted from yeast cells by Trizol and served as the template to obtain complementary DNA using the TransScript First-Strand cDNA Synthesis Kit (Trans, China). The converted cDNA and the specific primers was added to Top/Tip Green qRCR SuperMix to subject RT-PCR analysis employing the Roche LightCycler 96 Real-Time PCR System (Cal, US). ACT1 was selected as the internal reference gene.

### 2.4. Shake-flask cultivation

The engineered strains were grown at 42°C for high temperature stress or 1 g/L acetic acid for acetic acid stress in SD medium lacking uracil with 20 g/L glucose. Seed cultures were prepared by growing all strains individually in 15 mL culture tubes containing 2 mL medium at 30°C and 200 rpm overnight. Flasks (250 mL) containing 30 mL medium were then inoculated with the resulting seed cultures to an OD_600_ of 0.1. The strains were grown for 72 hours, and O_D600_ was measured every 12 hours using a spectrophotometer, model U-2900 (HITACHI, Chiyoda, Tokyo). All data represent the mean standard deviation from three independent experiments.

### 2.5. Cell viability analysis of engineered strains

A serial dilution assay was completed by taking samples at 24 h and serially diluting them 10-fold, followed by spotting 2.5 μL of the dilutions onto SD medium lacking uracil with 20 g/L glucose plates. The thermo tolerance and acetic acid tolerance ability of the engineered strains was characterized

### 2.6. ROS content determination

A total of 2×10^7^ cells were harvested by centrifugation, and ROS accumulation was determined by staining the cells with DCFH-DA, washing and diluting using PBS. In brief, cells were treated with 2 μM DCFH-DA dissolved in DMSO at 30°C for 15 min, then washed three times using PBS (pH 7.4) and tested for fluorescence intensity via a fluorescence micro-plate reader with the excitation wavelength of 488 nm and the emission wavelength of 525 nm (Cytation 3 Cell Imaging Multi-Mode Reader, Biotek, US).

## 3. Results and discussion

### 3.1. Analysis of GRXs in *S*.*cerevisiae*

To date, eight glutaredoxins (GRXs) have been characterized in Saccharomyces cerevisiae. These include dimeric GRXs (GRX1 and GRX2), multi-domain monothiol GRXs (GRX3 and GRX4), a mitochondrially targeted monothiol GRX (GRX5), and two monothiol GRXs (GRX6 and GRX7) associated with the Endoplasmic reticulum/Golgi apparatus. Additionally, GRX8, a disulfide GRX, is found in the cytoplasm (Fig. 1a).

**Fig. 1.**
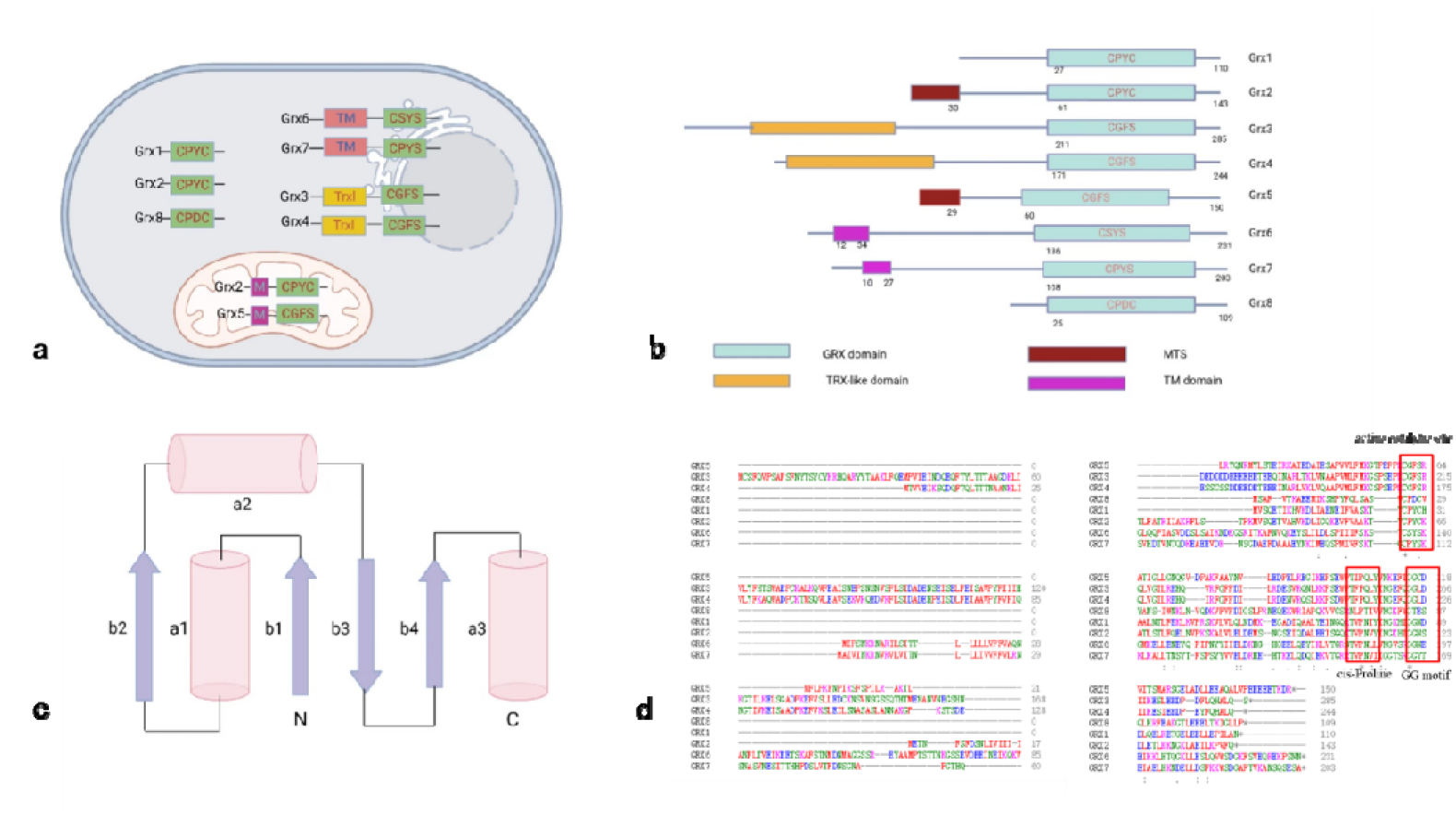
Domain structure of *S. cerevisiae* GRXs. (a) Species and distribution of GRXs (b) Structural domain (c) Secondary structure (d) Amino acid sequence alignment.

Structurally, GRXs are part of the thioredoxin (Trx) folding family. While prokaryotic GRXs typically possess a single domain, their eukaryotic counterparts often form multi-domain proteins (Fig. 1b). The GRX structure is predominantly composed of three alpha-helices surrounding four beta-sheets, featuring an active site similar to that of the Trx family oxidoreductases (Fig. 1c). This active site, often characterized by a Cys-X-X-Cys or Cys-XX-Ser motif, is situated within the flexible loop region between the first beta-sheet and alpha-helix. The N-terminal Cys residue of GRXs is exposed on the protein surface, akin to Trx family members. Adjacent to the active site are two critical regions: the cis-Proline-preceding amino acid (TVP) and the GRX-specific GG motif (GGxdD) (Fig. 1d). These motifs, in conjunction with GRXs and glutathione (GSH), facilitate the necessary catalytic reactions.

### 3.2. Construction of Engineered Strains

The Saccharomyces Genome Database (SGD,https://www.yeastgenome.org/) offers comprehensive genomic sequences for S. cerevisiae, including associated genes and gene products. In this study, we sourced eight GRX gene sequences from SGD and constructed a phylogenetic tree (Supplementary Fig. 1) to delineate their evolutionary relationships. Utilizing the S. cerevisiae strain BY4741 as a genomic template, we successfully amplified the eight GRX genes (Supplementary Fig. 2).

To achieve GRX overexpression, we employed the promoter TDH3p and terminator SLM5t to construct GRX devices within the expression vector POT1 (Fig. 2a, 2b). These vectors were transformed into the BY4741 strain, yielding eight distinct engineered strains, each overexpressing a unique GRX gene. A control strain, transformed with empty POT1 vectors, was also included.Reverse transcription-PCR (RT-PCR) data (Supplementary Fig. 3) confirmed upregulation of the GRX genes by 1.8 to 4.1-fold in the engineered strains relative to the control, validating the construction approach.

**Fig. 2.**
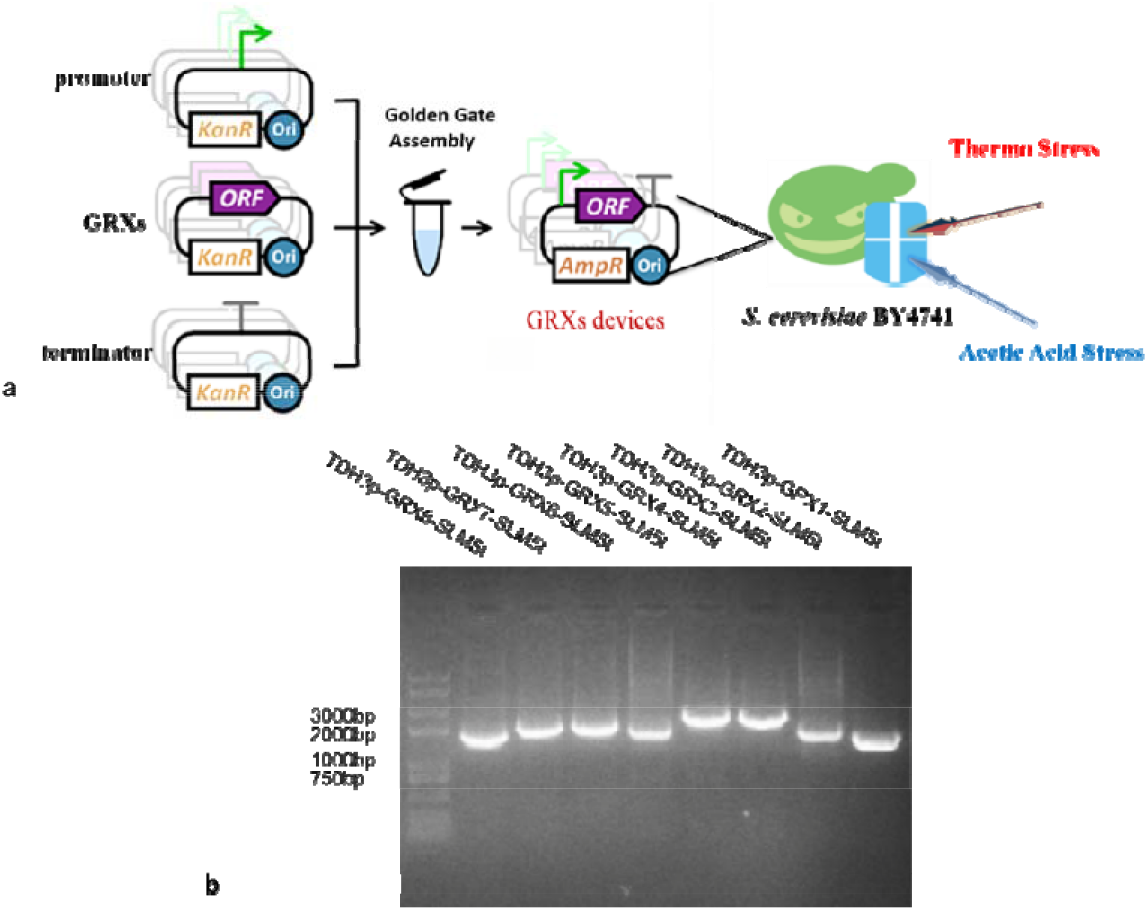
Construction of GRX devices using Golden Gate Assembly. (a) Overview of the construction strategy (b) Agarose gel electrophoresis of GRX devices.

### 3.3. Characterization of GRX devices in *S. cerevisiae*

The engineered strains were subjected to high-temperature stress (42□) or acetic acid stress (1 g/L) in a uracil-deficient SD medium. The growth curves revealed that strains overexpressing GRX1, GRX2, and GRX5 exhibited enhanced growth at 42□, whereas those overexpressing GRX3 showed inhibited growth (Fig. 3a). At 72 hours (Fig. 3b), the cell densities of the GRX1, GRX2, and GRX5 strains increased by 34.5%, 19.6%, and 27.96%, respectively, over the control strain. Correspondingly, cell viability was significantly improved (Fig. 3e). Under acetic acid stress (Fig. 3c), strains overexpressing GRX1, GRX2, GRX6, and GRX7 demonstrated superior growth, with the GRX8 overexpressing strain showing inhibited growth. At 72 hours (Fig. 3d), the cell densities of the GRX1, GRX2, GRX6, and GRX7 strains were 13.0%, 8.3%, 4.3%, and 5.9% higher than the control, respectively, with notable enhancements in cell viability (Fig. 3f).

**Fig. 3.**
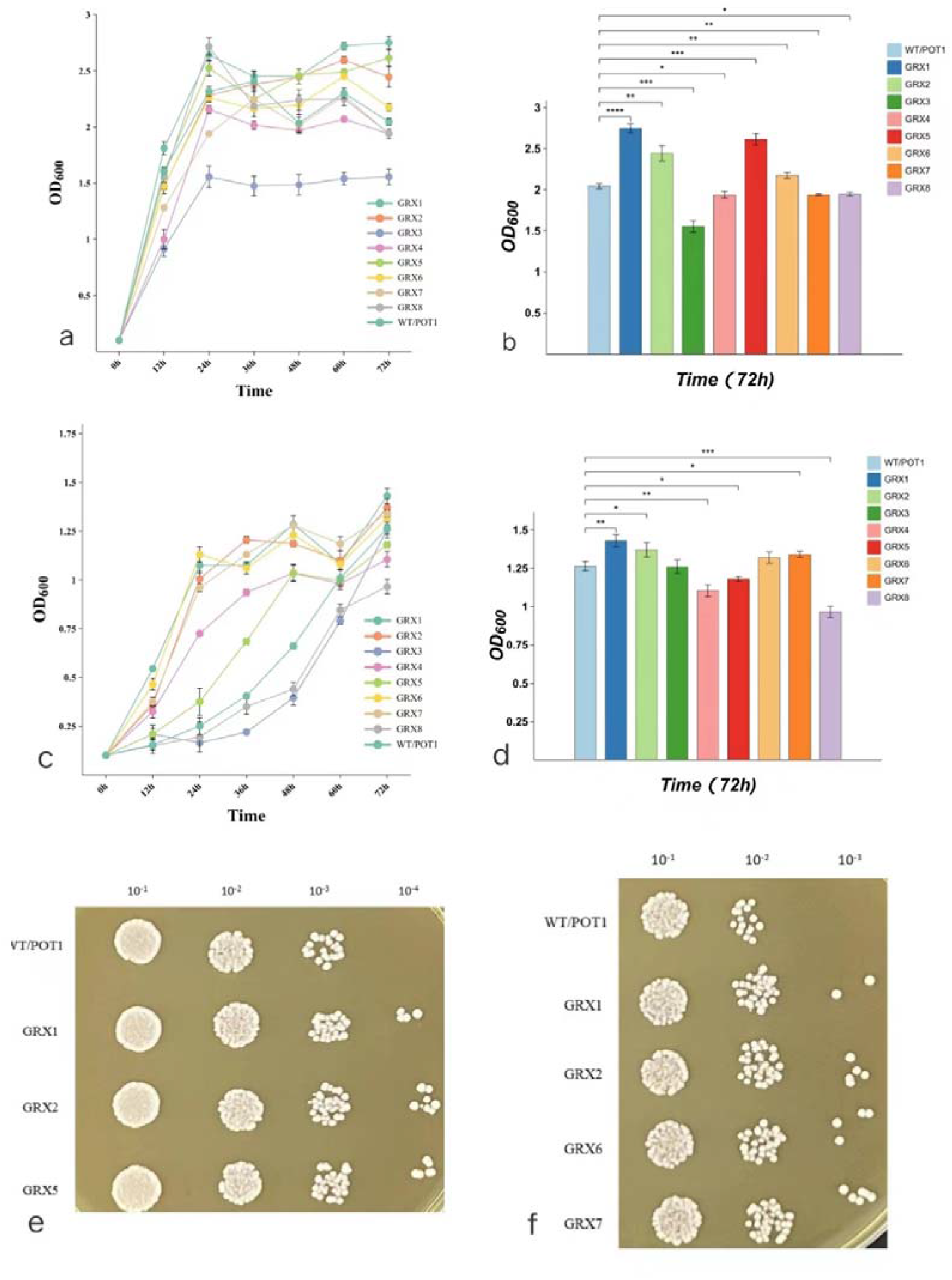
Cell growth and viability of engineered strains under stress conditions. (a) Growth curve at 42°C, (b) OD_600_ at 72 hours(42°C), (c) Growth curve under 1 g/L acetic acid stress, (d) OD_600_ at 72 hours(1 g/L acetic acid), (e) Cell viability at 72 hours(42°C), (f) Cell viability at 72 hours(1 g/L acetic acid).

### 3.4. Detection and analysis of ROS level in engineered strans

Elevated temperatures can denature proteins in the electron transport chain of S. cerevisiae, leading to electron leakage and accumulation of reactive oxygen species (ROS). The fluorescence probe DCFH-DA is utilized for ROS detection. Non-fluorescent DCFH-DA enters the cell and is hydrolyzed by esterases to DCFH, which remains cell-trapped. Intracellular ROS oxidizes DCFH to fluorescent DCF, allowing ROS levels to be quantified via DCF fluorescence intensity.

Our findings indicate that strains overexpressing GRX1, GRX2, and GRX5 displayed enhanced growth at 42°C. GRX1 and GRX2, which share 64% sequence similarity, are primarily cytoplasmic, with GRX2 also present in mitochondria. They serve as disulfide oxidoreductases and glutathione transferases (GSTs), facilitating glutathione complex formation and detoxification. GRX5, located in the mitochondrial matrix, is involved in Fe/S center synthesis and assembly, influencing ROS production. A knockout of GRX5 results in increased ROS levels, highlighting its role in antioxidant defense.

The ROS detection results (Fig. 4) show that the engineered strains overexpressing GRX1, GRX2, and GRX5 had significantly lower intracellular ROS levels than the control strain under high-temperature conditions. After 24 hours at 42°C, the ROS content in these strains was 36.2%, 41.2%, and 58.8% that of the control, respectively, suggesting the GRX overexpression effectively mitigated ROS accumulation.

**Fig. 4.**
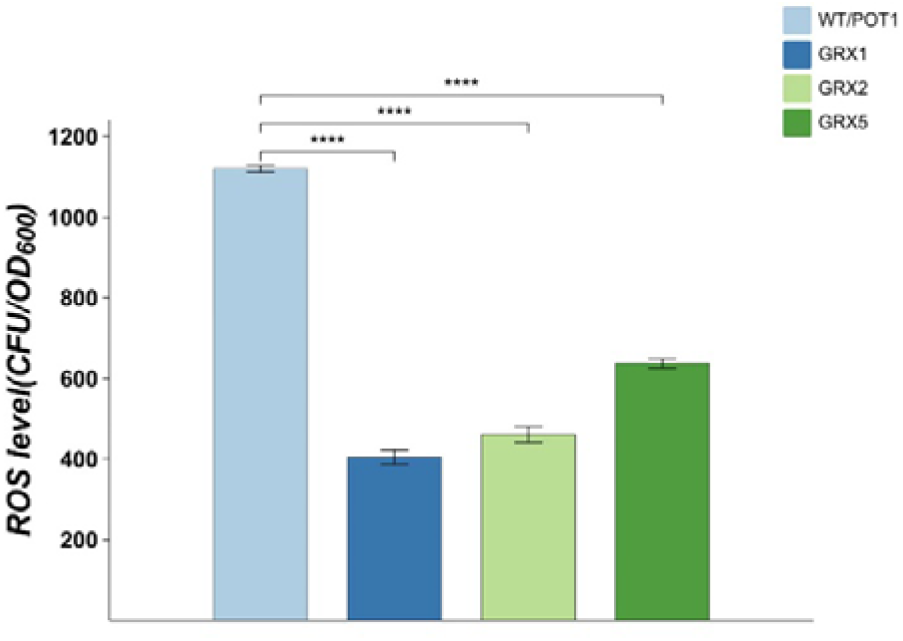
ROS content detection in engineered strains.

## 4. Conclusion

Through the application of engineering principles and synthetic biology techniques, we have successfully enhanced the stress resilience of microbial cell factories, specifically S. cerevisiae, under challenging conditions. Our approach, which involves the introduction of a GRX device to bolster the yeast”s tolerance to both thermal and acetic acid stress, presents a novel strategy for strain improvement. Although the precise mechanisms underlying the GRX device”s enhancement of acid tolerance in S. cerevisiae are not fully elucidated, our results unequivocally demonstrate its efficacy in improving yeast resistance to the specified stresses. This advancement offers a promising avenue for increasing the efficiency of biomanufacturing processes and markedly reducing energy usage, a concept that holds potential for broader application across various microbial strains.

Looking ahead, a more profound comprehension of microbial metabolic regulation and the molecular underpinnings of stress resistance will empower us to design sophisticated, multi-tiered defense systems. Such an integrated approach will not only refine the stress resilience of cell factories but also adapt them to thrive in a spectrum of hostile environments. The anticipated outcomes include significant advancements in the productivity of biological manufacturing operations, alongside a reduction in the energy footprint and overall production costs associated with fermentation processes.

## Author statement

Lin Wang: Methodology, Construction and fermentation validation of engineered strains, Writing – review & editing. Li-shuang Zhang: Construction and fermentation validation of engineered strains, Writing – review & editing. Mei-ling Zhang: Methodology, Construction and fermentation validation of engineered strains, Writing – review & editing. Ya-xin He: Methodology, Fermentation experiments. Yuan Yu: Methodology, Supervision. Ke Xu: Methodology, Writing – review & editing.

## Acknowledgements

This work was financially supported by the Natural Science Foundation of China (No.22078171, No. 32171430), the S&T Program of Hebei (21374301D), the Natural Science Foundation of Hebei Province (No. B2021209008) and the S&T Program of Tangshan (21130201C).

